# Translational accuracy of a tethered ribosome

**DOI:** 10.1101/2020.03.12.988394

**Authors:** Celine Fabret, Olivier Namy

**Affiliations:** Université Paris-Saclay, CEA, CNRS, Institute for Integrative Biology of the Cell (I2BC), 91198, Gif-sur-Yvette, France

## Abstract

Ribosomes are evolutionary conserved ribonucleoprotein complexes that function as two separate subunits in all kingdoms. During translation initiation, the two subunits assemble to form the mature ribosome, which is responsible for translating the messenger RNA. When the ribosome reaches a stop codon, release factors promote translation termination and peptide release, and recycling factors then dissociate the two subunits, ready for use in a new round of translation. A tethered ribosome, called Ribo-T, in which the two subunits are covalently linked to form a single entity, was recently described in *Escherichia coli*^1^. A hybrid ribosomal RNA (rRNA) consisting of both the small and large subunit rRNA sequences was engineered. The ribosome with inseparable subunits generated in this way was shown to be functional and to sustain cell growth. Here, we investigated the translational properties of Ribo-T. We analyzed its behavior in −1 or +1 frameshifting, stop codon readthrough, and internal translation initiation. Our data indicate that covalent attachment of the two subunits modifies the properties of the ribosome, altering its ability to initiate and terminate translation correctly.

## Introduction

Translation is the last step in gene expression, in which the coding sequence of the messenger RNA (mRNA) is translated into the amino-acid sequence of the corresponding protein. Ribosomes catalyze protein synthesis, a vital cellular activity. Translation is a highly dynamic process, with four major phases: initiation, elongation, termination and ribosome recycling. During each phase, ribosomes form transient complexes with auxiliary translation factors that facilitate protein synthesis^2^.

In all kingdoms of life, ribosomes consist of two subunits. The 30S subunit contains the 16S rRNA and 21 ribosomal proteins responsible for decoding genetic sequences. The 16S rRNA is involved in recognition of the Shine-Dalgarno (SD) sequence or ribosome binding site of the mRNA when it emerges from the RNA polymerase. This sequence is complementary to the 3’end of the 16S ribosomal RNA (anti-Shine-Dalgarno (ASD) sequence). The 50S subunit contains two rRNA molecules, the 5S and 23S rRNAs, with 33 proteins. It is responsible for catalyzing peptide bond formation. These two subunits associate with each other during translation initiation, rotate during elongation, and dissociate after protein release. This organization is thought to be essential for biogenesis, successful protein synthesis and cell viability.

A ribosome with tethered subunits, in which the two subunits were covalently linked in a single entity, was recently engineered in *Escherichia coli*^1,3^. The hybrid ribosomal RNA (rRNA) consisted of the small and large subunit rRNA sequences, linked by short RNA linkers. The resulting ribosome, Ribo-T, with its inseparable subunits, was shown to be functional and able to sustain the cell growth, although doubling times were slower (107 min for Ribo-T, 35 min for wild-type ribosomes). The rate protein synthesis with Ribo-T catalysis was 50% that with wild-type ribosomes. The slow growth of Ribo-T cells was probably due to Ribo-T biogenesis being slower than the biogenesis of wild-type ribosomes, rather than impaired Ribo-T activity^3^.

Cell viability depends on a balance between rapid protein synthesis and accurate decoding of the genetic information. Translational error rates remain low throughout all steps in the process^4^. However, signals present in specific mRNAs at defined locations can program a high rate of errors^5^. These recoding events can occur during translation elongation (frameshifting) or at the termination step (stop codon readthrough). During frameshifts, the ribosome is induced to shift to an alternative, overlapping reading frame, whereas, in stop codon readthrough, the specific context of the termination codon may promote decoding with a near-cognate tRNA rather than a release factor. These sequences have specific impacts on translation fidelity, and are therefore powerful tools for investigations of the translation accuracy of Ribo-T ribosomes.

In this study, we investigated the translational properties of Ribo-T ribosomes. Do these ribosomes maintain the correct reading frame as faithfully as wild-type ribosomes? Do they terminate translation accurately at stop codons? Do they translate messenger RNA accurately? We investigated the behavior of Ribo-T ribosomes with recoding signals instructing the ribosome to change translational reading frame (frameshifting) or to read stop codons as sense codons (stop codon readthrough). We also investigated the preferential mode of translation used by Ribo-T ribosomes to express adjacent open reading frames on the same mRNA, given that the two subunits were inseparable. Our data reveal an impairment of tethered ribosomes for +1 frameshifting and termination, but show that these ribosomes are otherwise as effective as wild-type ribosomes.

## Materials and Methods

### *Escherichia coli* strains

The experiments were performed with the *E. coli* BL21 (DE3) strain (Invitrogen), with and without the poRibo-T2 vector carrying the Ribo-T ribosomal DNA for the expression of Ribo-T ribosomes^1^. The ASD sequence of the Ribo-T 16S rRNA was altered from the wild-type sequence (5’-UCACCUCCUUA-3’) to an orthogonal sequence (5’-UCAUUGUGGUA-3’).

### Plasmids

We used the bacterial pCL99 dual reporter system. This plasmid carries the *lacZ-luc* fusion gene, encoding the β-galactosidase and luciferase enzymes, under the control of the T7 promoter, together with the streptomycin resistance gene, and the CloDF13-derived CDF replicon^6^. The various target sequences tested for translation accuracy were inserted by oligo-cloning into the *Msc*I restriction site of pCL99, which placed them at the junction between the *lacZ* and *luc* genes. The cloned double-stranded oligonucleotides were obtained by hybridizing the two complementary oligonucleotides in ligation buffer after heating for 5 min at 100°C and incubation at room temperature. In-frame controls were constructed for each target sequence. All constructs were verified by sequencing. The target sequences are listed in Table 1.

**Table 1:**
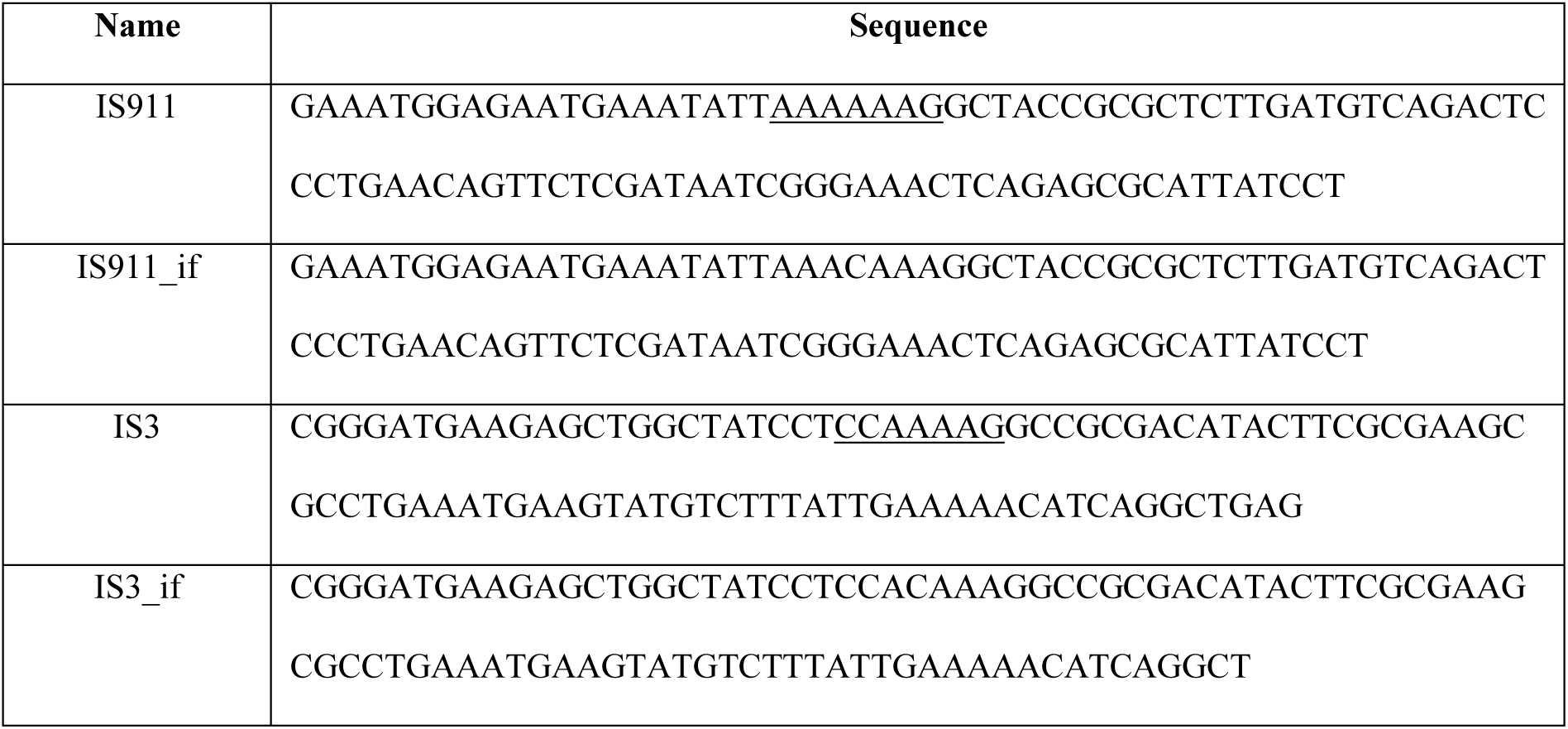
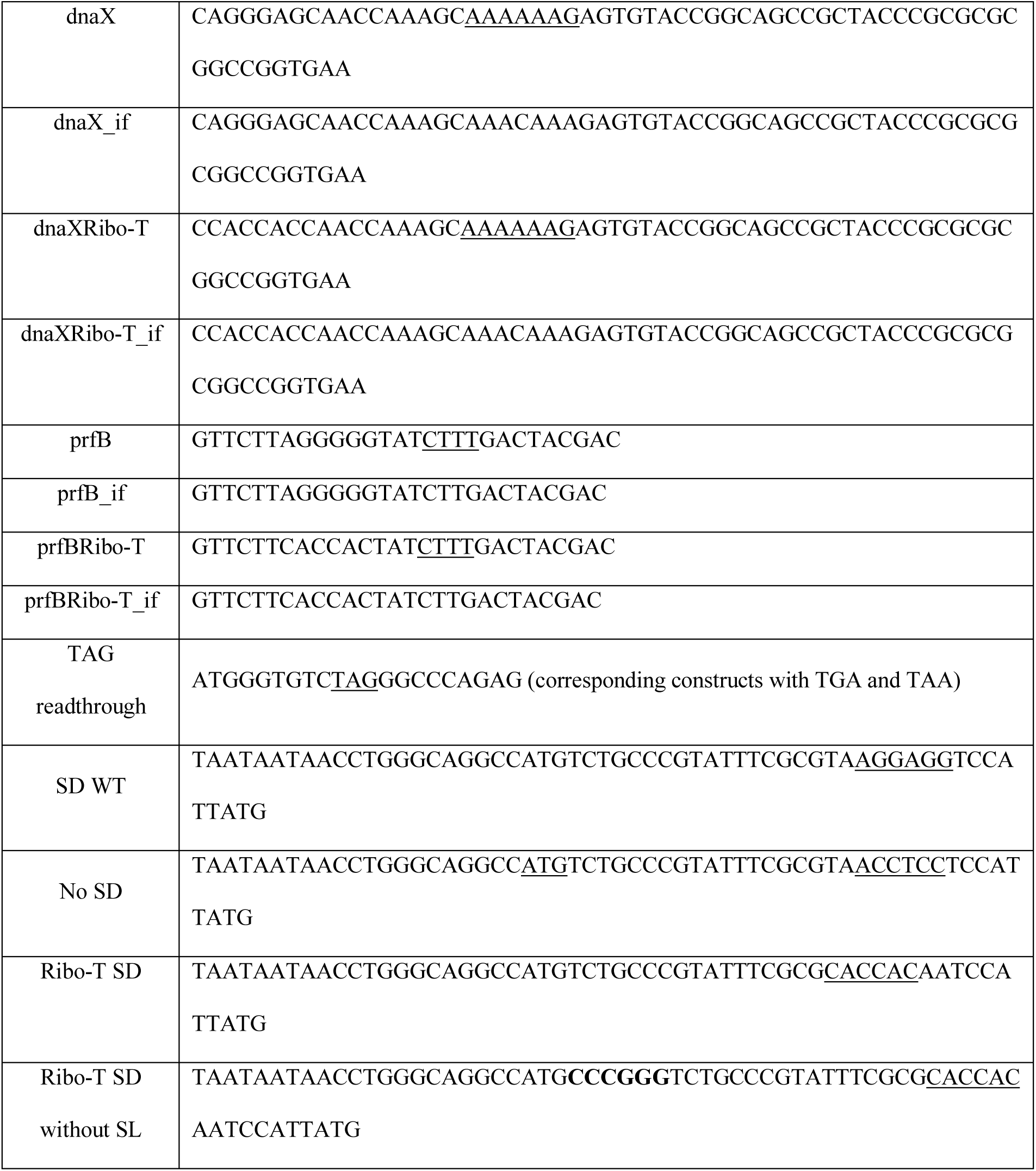
Target sequences used in the study. For frameshifting sites, the slippery sequence is underlined. In-frame control sequences corresponding to each target sequence are indicated by “_if”. For −1 frameshifting, a C was added within the slippery sequence; for +1 frameshifting, the T of the TGA stop codon was removed; and for stop codon readthrough, the stop codon was replaced by AAG. The *dnaX* and *prfB* frameshifting sites were also obtained with a specific CACCAC Ribo-T SD sequence upstream from the slippery sequence (dnaXRibo-T and prfBRibo-T, respectively). The *Sma*I site used for stem-loop (SL) cloning is indicated in bold typeface.

For creation of the pCL99_Ribo-T constructs, containing the *lacZ* gene controlled by an orthogonal SD sequence, the wild-type SD sequence (AGGAGG) was mutated to an orthogonal sequence, CACCAC, recognized by the Ribo-T ribosomes. A PCR fragment encompassing the *lacZ* SD sequence from the *Acl*I to *Fsp*I sites of the vector was generated by PCR with the mutated sequence. An upstream PCR fragment was amplified with the oligonucleotides 5’-CTCTGCGACATCGTAT<u>AACGTT</u>ACTGG-3’ (*Acl*I site underlined) and 5’-GACCGTAATCATGGTATATT<u>GTGGTG</u>TTAAAGTTAAACAAAATTATTTC-3’ (mutated SD sequence underlined), and a downstream PCR fragment was amplified with the oligonucleotides 5’-GTTTAACTTTAA<u>CACCAC</u>AATATACCATGATTACGGAC-3’ (mutated SD underlined) and 5’-CGCCATTCAGGC<u>TGCGCA</u>ACTGTTGGG-3’ (*FspI* site underlined). These two PCR fragments were joined by PCR with equimolar amounts of the two fragments as the template and oligonucleotides containing the *Acl*I and *Fsp*I sites. The resulting PCR fragment was digested with *Acl*I and *Fsp*I and inserted into the pCL99 constructs predigested with the same enzymes. The replacement of the wild-type SD sequence with the mutated sequence was verified by sequencing, for all pCL99_Ribo-T constructs.

For constructs with the intergenic sequence containing a stem-loop structure (SL), the 5’-GCGATATCCCGTGGAGGGGCGCGTGGTGGCGGTGCT-3’ and the phosphorylated 5’-CACCACGCGCCCCTCCACGGGATATCGC-3’ oligonucleotides were hybridized, as were the phosphorylated 5’-GCAGCACCGCCACCACGCGCCCCTCCACGGGATATCGCT-3’ and 5’-AGCGATATCCCGTGGAGGGGCGCGTGGTGGCGGTGCTGCAGCACCGC-3’ oligonucleotides. These two pairs of annealed oligonucleotides were then ligated together to produce the 75 bp fragment forming a −105 kcal/mol stem-loop structure (from RNA folder WebServer). This fragment was inserted into the *Sma*I site of the pCL99_Ribo-T Ribo-T SD without SL. The presence of the correct SL was verified by sequencing.

### Quantification of accurate target sequence translation

*E. coli* BL21 or BL21 poRIBO-T2 competent cells were transformed with the pCL99 or pCL99_Ribo-T constructs, respectively, carrying the various target sequences used to test the translation accuracy of ribosomes. The transformants were cultured overnight at 30°C in 500 µl of Luria broth supplemented with the appropriate antibiotics (50 µg/ml streptomycin, 100 µg/ml ampicillin). Cultures were centrifuged for 2 min at room temperature, and the cell pellets were resuspended in 100 µl of luc buffer (25 mM Tris-phosphate pH 7.8, 8 mM MgCl_2_, 1 mM DTT, 1 mM EDTA, 1% Triton X-100, 0.1% BSA, 15% glycerol and cOmplete Protease Inhibitor Cocktail (Roche)) with 50 µl of acid-washed glass beads (Sigma)^7^. Cells were lysed by vortexing for 30 min at 4°C. Luciferase and β-galactosidase activities were quantified, in 1 µl and 20 µl of cell extract, respectively^7^.

The translational efficiency of the ribosomes was calculated as the ratio of luciferase activity to β-galactosidase activity for the target, divided by the ratio of luciferase activity to β-galactosidase activity for the corresponding in-frame construct. We calculated the median value for at least six independent experiments. The significance of differences was determined in Student’s *t-*tests.

We used constructs with a *lacZ* gene carrying a wild-type SD sequence AGGAGG to analyze the translational efficiency of wild-type ribosomes. For assessments of the translational efficiency of Ribo-T ribosomes, we used the corresponding constructs with a mutated SD sequence CACCAC upstream from the *lacZ* gene, in the presence of the vector encoding the Ribo-T ribosomes. The mutated SD sequence was complementary to the 3’ end of the 16S ribosomal RNA of the Ribo-T ribosomes. In strains containing this vector, both wild-type and Ribo-T ribosomes are present, but only the Ribo-T ribosomes can initiate *lacZ* translation, because of the mutated SD sequence. Crosstalk between wild-type ribosomes and the orthogonal mRNA has recently been described^8^. We checked the level of crosstalk in our system, and found that it was too low to affect the results. Indeed, in the presence of wild-type ribosomes only, constructs with a mutated SD sequence upstream from the *lacZ* gene generate too little β-galactosidase activity for measurement in our conditions. Similarly, expression of the *luc* gene with the mutated SD sequence resulted in levels of activity one tenth those observed with Ribo-T ribosomes (see also below).

## Results

We first investigated the way in which Ribo-T ribosomes deal with programmed frameshifting sequences. In essence, frameshifting is triggered by two features: a slippery sequence favoring tRNA slippage and one or several stimulatory elements that enhance the process by inducing a ribosomal pause.

### Ribo-T ribosomes are prone to frameshift independently of the SD sequence

We selected three programmed −1 frameshifting sites from the IS911 and IS3 insertion sequence elements and the *E. coli dnaX* gene (Figure 1)^9^. Frameshifting is stimulated by two elements: an RNA structure (a stem-loop for IS911 and *dnaX*, a pseudoknot for IS3) located downstream from the slippery site, and an SD sequence located upstream from the slippery site. The combination of these elements, together with the highly efficient slippery sequence, gave a −1 frameshifting efficiency of about 13%, 8% and 60% for IS911, IS3 and dnaX, respectively^10-12^.

**Figure 1:**
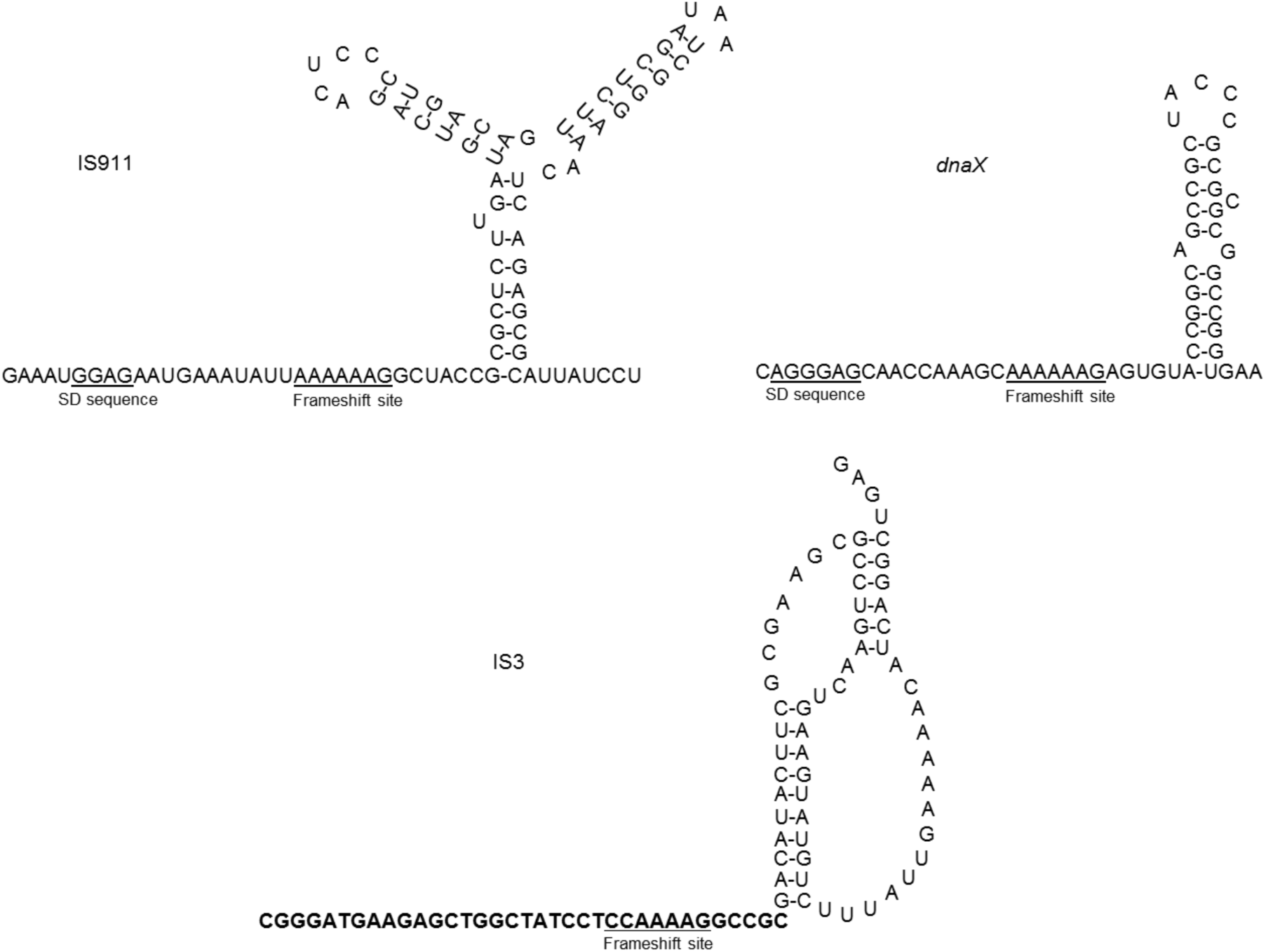
The −1 frameshifting sites.

We first measured frameshifting efficiencies with WT sequences (i.e. with a WT SD). As Ribo-T ribosomes cannot base pair to the WT SD, we assumed that they would be unable to frameshift. Surprisingly, this turned out not to be the case, as the efficiency of −1 ribosomal frameshifting was 13% for IS911 with wild-type ribosomes and 14% with Ribo-T ribosomes (*p*-value of 0.54), 33% for IS3 with either type of ribosome (*p*-value of 0.77), 43% for *dnaX* with wild-type ribosomes and 45% with Ribo-T ribosomes (*p*-value of 0.77) (Figure 2A). These results suggest that Ribo-T ribosomes frameshift independently of the SD. We investigated this possibility by performing the reverse experiment. We replaced the *dnaX* SD sequence with an SD recognized by Ribo-T ribosomes (dnaXRibo-T). This modification greatly decreased the frameshifting efficiency of WT ribosomes (11% versus 43%; *p*-value of 1.7×10^−3^), but this change had no significant impact on Ribo-T ribosome frameshifting (45% vs. 44%; *p*-value 0.85) (Figure 2B), confirming that Ribo-T ribosomes are insensitive to the presence of an SD sequence even though they maintain high levels of −1 frameshifting.

**Figure 2:**
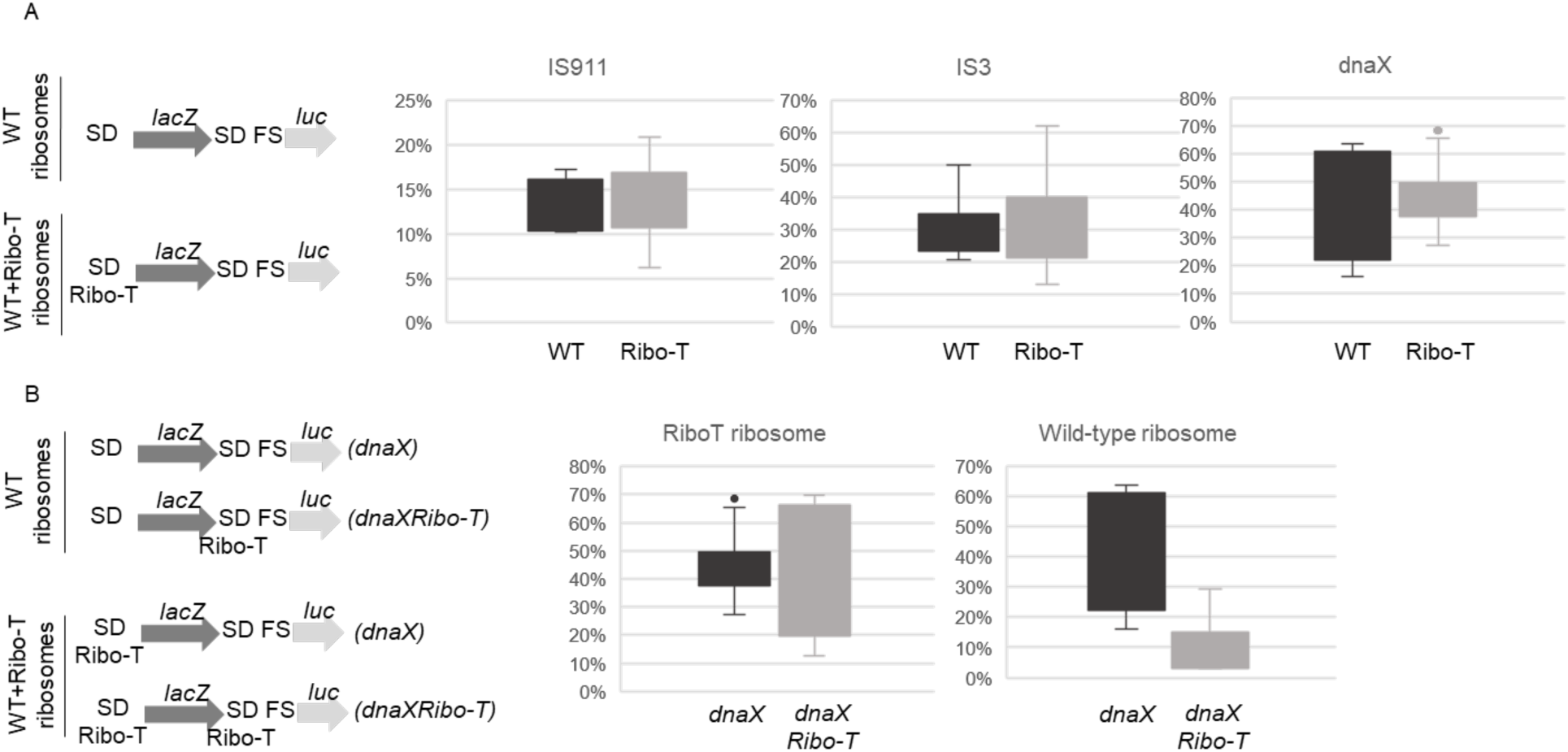
Translational efficiencies during −1 frameshifting. The constructs analyzed are shown on the left, with the type of ribosomes present in the cells. The percent frameshifting is shown for the IS911, IS3 and *dnaX* sequences translated by wild-type (WT) (dark gray) and Ribo-T (light gray) ribosomes (A); and for the *dnaX* (dark gray) and *dnaXRibo-T* (light gray) sequences translated by Ribo-T or wild-type ribosomes (B). The UGA stop codon in the −1 frame of the *dnaX* frameshifting site was changed to an UGU codon for translation of the *luc* gene after frameshifting.

This surprising result led us to investigate whether this effect was specific to −1 frameshifting signals, or also applied to +1 frameshifting.

For +1 frameshifting, we selected the site from the *E. coli prfB* gene (Figure 3A). Frameshifting efficiency is typically around 30% - 50%, and this process is dependent on a poor termination context of the stop codon (UGA C), together with the stimulatory effect of an SD sequence located upstream from the slippery site^13^.

**Figure 3:**
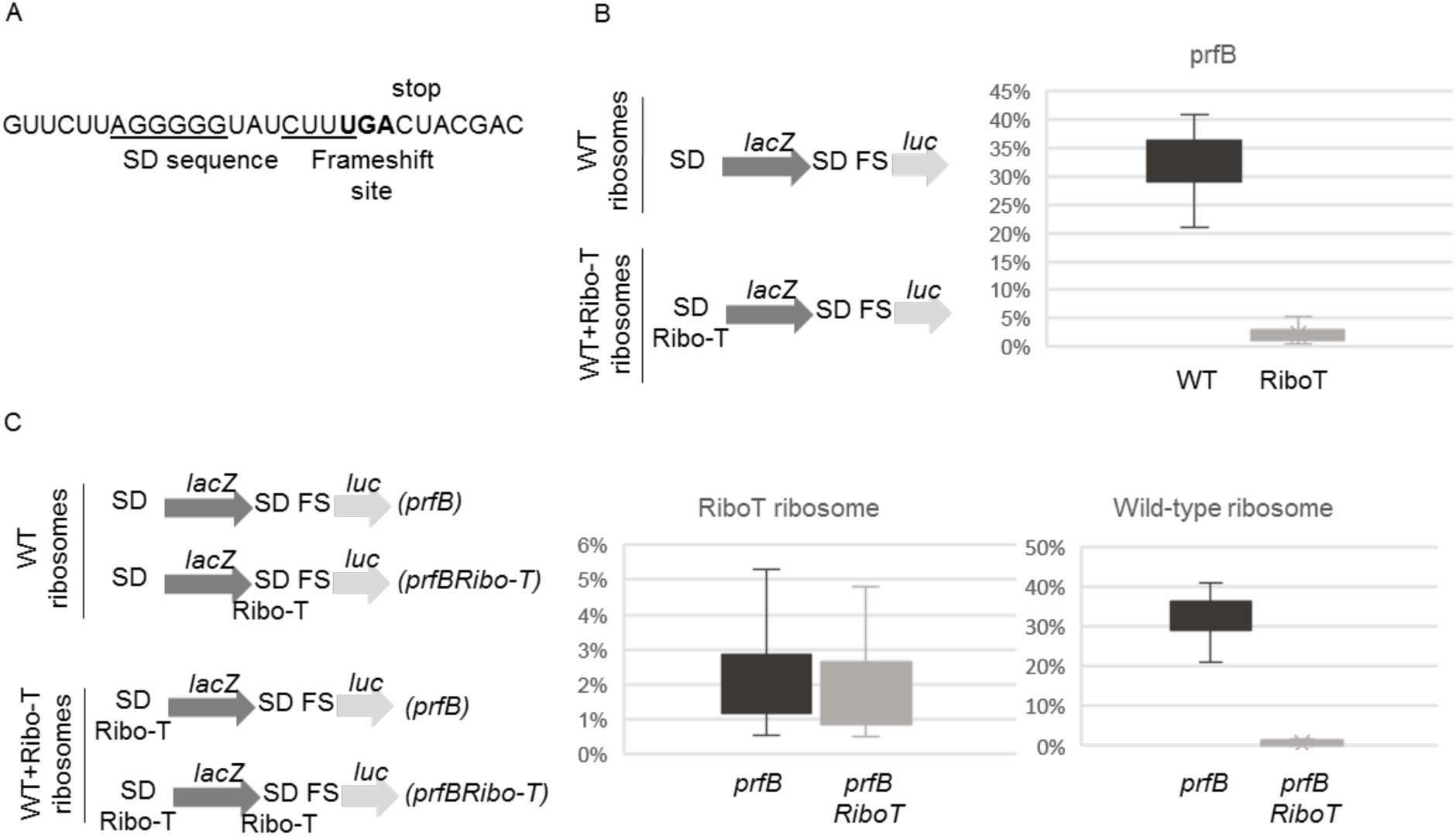
*prfB* +1 frameshifting. Sequence of the *E. coli* frameshifting site (A). Translational efficiencies for *prfB* frameshifting, for wild-type ribosomes (dark gray) and Ribo-T ribosomes (light gray) (B); or for the *prfB* (dark gray) and *prfBRibo-T* (light gray) sequences translated by Ribo-T or wild-type ribosomes (C). The constructs analyzed are shown on the left, with the type of ribosomes present in the cells.

We obtained a +1 frameshifting efficiency of 33% for the WT ribosome, confirming previous results^14^. However, Ribo-T ribosomes frameshift inefficiently on this sequence, with an efficiency of only 2% (*p*-value of 8.9×10^−9^ versus wild-type ribosomes; Figure 3B). This may be because the stimulatory SD sequence is wild-type and is not, therefore, recognized by Ribo-T ribosomes. We addressed this possibility, by replacing the wild-type SD sequence with a sequence recognized by Ribo-T ribosomes. Interestingly, this did not restore the +1 frameshifting efficiency of Ribo-T ribosomes (1.8%, *p*-value of 0.56 relative to the previous finding), but it did strongly decrease the +1 frameshifting activity of wild-type ribosomes (0.6% versus 33%, *p*-value of 1.6×10^−5^), which were unable to recognize this sequence as an SD sequence (Figure 3C).

These results confirm that Ribo-T ribosomes are unable to sense SD sequences and use them as stimulatory elements for frameshifting to either the −1 or +1 frame.

### Ribo-T ribosomes make inefficient use of the SD sequence to initiate translation

Our data demonstrate the inability of Ribo-T ribosomes to use SD sequences to stimulate frameshifting. This led us to investigate the recognition of the SD sequence during translation initiation, when this sequence is used to assemble the ribosome at the start codon. We used the 46-nucleotide intergenic *E. coli lacZ-lacY* sequence, which carries the SD sequence promoting initiation at the *lacY* start codon to address this question. We modified this sequence by deleting the SD sequence or replacing it with the Ribo-T SD sequence recognized only by Ribo-T ribosomes (see Materials and Methods, Table 1). In this system, *lacZ* could be translated only by wild-type ribosomes, because all constructs carried a wild-type SD sequence upstream from *lacZ*. The ability of wild-type or Ribo-T ribosomes to translate *luc* depended on whether the second SD sequence was recognized.

In the absence of Ribo-T, we observed almost no luciferase activity when the SD sequence upstream from the *luc* gene was removed. By contrast, when the wild-type SD sequence was used, we obtained high levels of luciferase activity, demonstrating the correct reconstitution of the SD dependence of translation initiation context by the reporter system (Figure 4A). In the presence of the Ribo-T SD sequence, we observed no statistically significant difference (*p*-value 0.04, versus no SD) in luciferase activity, which suggests that wild-type ribosomes cannot use this alternative SD sequence to initiate translation (Figure 4B). Interestingly, in the presence of Ribo-T ribosomes, luciferase induction was increased by a factor of four (*p*-value 4.7×10^−4^), indicating that Ribo-T ribosomes can use this alternative SD sequence, albeit inefficiently relative to WT ribosomes in the presence of a wild-type SD sequence (Figure 4B).

**Figure 4:**
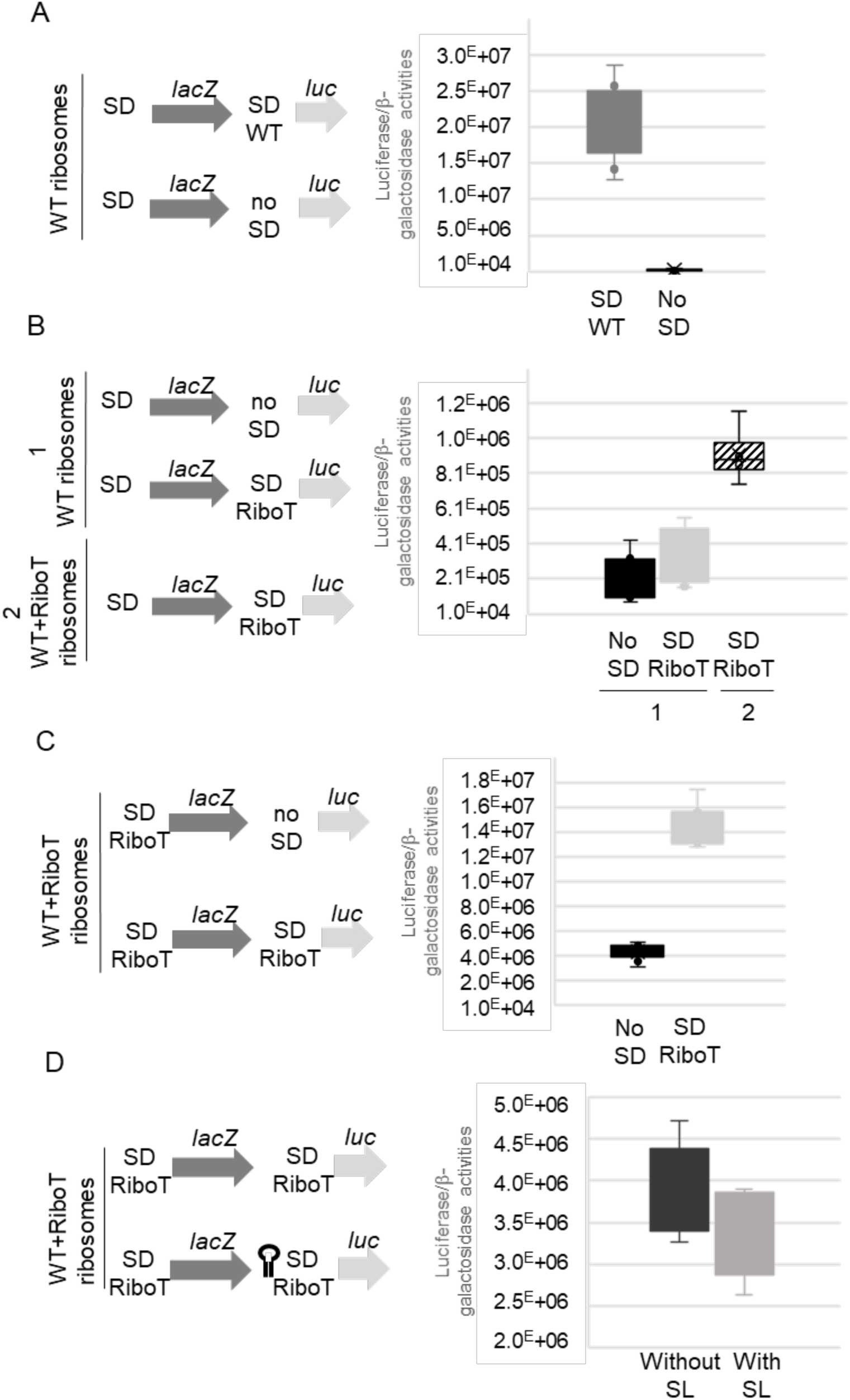
Internal initiation of wild-type ribosomes. The constructs analyzed are shown on the left, with the type of ribosomes present in the cells. The ratio of luciferase activity to β-galactosidase activity measured for each construct is shown in the right box plot. Translations of the *lacZ* gene by wild-type ribosomes (WT), and of the *luc* gene with either a wild-type SD (WT SD) (dark gray) or no SD (black) by wild-type ribosomes (A). Translations of *lacZ* by wild-type ribosomes, and of *luc* with either no SD (black) or a Ribo-T SD (light gray) by wild-type ribosomes (1), or a Ribo-T SD (hatched) by wild-type and Ribo-T ribosomes (2) (B). Translations of *lacZ* by Ribo-T ribosomes and of *luc* with either no SD (black) or a Ribo-T SD (light gray) by wild-type and Ribo-T ribosomes (C), or by Ribo-T ribosomes with (light gray) or without (black) a stem-loop structure upstream from *luc* (D).

In prokaryotes, ribosomes are thought to initiate the translation of a downstream gene either by 70S re-initiation after translation of the upstream gene is terminated, or by internal loading of the 30S at the initiation site (70S for Ribo-T with tethered subunits)^15,16^. Both these processes depend on the distance between the upstream stop codon and the downstream start codon, with a seven-codon interval thought to be crucial for the posttermination reinitiation of protein synthesis by ribosomes. The long intergenic sequence between *lacZ* and *luc*, with three successive stop codons ending the first ORF, would not favor a scanning process.

We investigated whether the type of ribosomes translating *lacZ* influenced the rate of translation initiation for the *luc* gene, by replacing the wild-type SD sequence upstream from *lacZ* with a Ribo-T SD sequence. We reasoned that if 70S re-initiation could occur, we would observe an increase in luciferase activity due to Ribo-T ribosomes initiating translation upstream in *lacZ*. We found that expression of the luciferase reporter gene was four times stronger when the Ribo-T SD sequence was placed in front of the start codon for the upstream *lacZ* than in the total absence of an SD sequence (no-SD; *p*-value 9.8×10^−6^; Figure 4C). This increase was of a similar magnitude to that obtained when a wild-type SD sequence was placed upstream from the *lacZ* gene (Figure 4B). The higher activity ratio obtained for the Ribo-T ribosomes translating both *lacZ* and *luc* (Figure 4C) resulted from lower levels of β-galactosidase activity than were obtained when wild-type ribosomes translated *lacZ* (SD Ribo-T data in figure 4B(2)). Thus, translation initiation at the *luc* gene is not dependent on the nature of the ribosomes translating the upstream *lacZ* gene. For confirmation of this observation, we inserted a stable stem loop structure immediately downstream from *lacZ*, to act as a roadblock for potential scanning ribosomes. We found that structure had no effect on the initiation of *luc* translation (*p*-value 0.22), strongly suggesting that the initiation of *luc* translation occurred through the internal loading of Ribo-T ribosomes (Figure 4D). Overall, our results indicate that Ribo-T ribosomes do not use SD sequence efficiently during either initiation or programmed frameshifting.

### Ribo-T ribosomes terminate translation less efficiently at stop codons

Stop codon readthrough occurs when the ribosome reads the termination codon as a sense codon and continues translation until the next stop codon in the same reading frame^17^. Quantifying stop codon readthrough is an efficient way to assess the accuracy of ribosomes and their ability to accommodate near-cognate tRNAs. Unfortunately, no natural stop codon readthrough sequence has ever been identified in *E. coli*. We therefore used the stop codon sequence of the *mtmB1* gene encoding monomethylamine methyltransferase from the archaea *Methanosarcina barkeri*^18^, which allows the insertion of a pyrrolysine, to estimate the accuracy of Ribo-T ribosomes during translation termination. This sequence has been shown to promote stop codon readthrough in *E. coli*, even in the absence of pyrrolysine^6^.

We found that Ribo-T ribosomes terminated translation less efficiently than wild-type ribosomes (Figure 5; 0.07% vs. 0.013%, *p*-value of 4.7×10^−7^; 0.26% vs. 0.11%, *p*-value of 5×10^−3^; 0.04% vs. 0.01%, *p-*value of 2.5×10^−4^, for UAG, UGA and UAA stop codons, respectively). Thus, Ribo-T ribosomes display a slight impairment of stop codon recognition.

**Figure 5:**
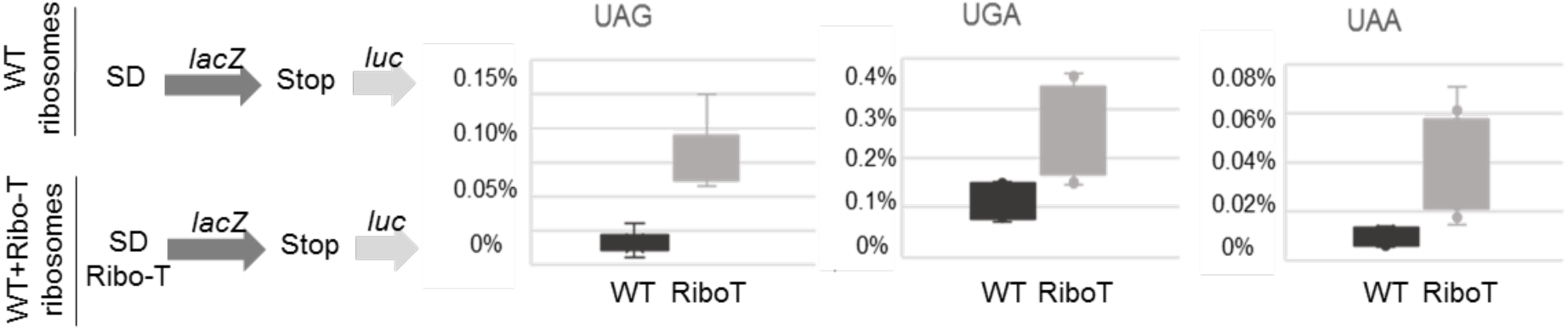
Translational readthrough efficiencies. The percent readthrough is shown for them three stop codons encountered by the wild-type (WT) (dark gray) and Ribo-T (light gray) ribosomes. The constructs analyzed are shown on the left, with the type of ribosome present in the cells.

## Discussion

In all organisms, the ribosome consists of two unequal subunits, the large and small ribosomal subunits, which associate with each other in a labile manner, via bridges. It remains unclear why the ribosome consists of two loosely associated subunits that must be associated for translocation to occur. It has been suggested that the mutual mobility of the two ribosomal subunits is essential for the translocation mechanism. However, a functional ribosome with covalently linked subunits has been successfully created independently by two groups^1,19^, calling this hypothesis into question, or suggesting that the covalently linked structure retained some mobility.

Surprisingly, tethered ribosomes (also called Ribo-T ribosomes) support cell growth and are not associated with any obvious phenotype, other than a slow-growth phenotype that has been attributed to slower rRNA processing rather than translational defects^3^. We investigated the translational accuracy of Ribo-T ribosomes. For this purpose, we constructed reporters for measuring translational errors at stop codons and in reading frame maintenance (frameshifting). Such recoding events constitute useful tools for studying translation accuracy, as the sequences concerned alter ribosome proofreading activity, and translational defects are amplified at these sites. To our surprise, we observed no change in −1 frameshifting efficiency, whatever the type of SD sequence (either wild-type or Ribo-T) used (Figure 2B). These initial results were confirmed for +1 frameshifting (Figure 3C). Interestingly, the efficiency of Ribo-T ribosomes was much lower for +1 frameshifting than for −1 frameshifting (Figures 2A and 3B). This may reflect the different roles of SD-ASD interaction in these two events. Indeed, for +1 frameshifting, this interaction causes the release of the deacylated E-site tRNA, destabilizing the ribosome complex and affecting P-site tRNA slippage^20,21^. Both types of frameshifting are known to involve base pairing between the SD sequence and the ASD sequence on the 16S rRNA to increase frameshifting efficiency, but the mechanisms of these two types of frameshifting event are different^20,22-25^.

These results indicate that Ribo-T ribosomes do not use SD sequences to promote frameshifting. This situation is atypical, because frameshifting occurs during the elongation phase and SD sequences are used at the initiation step. This may reflect a general trend towards Ribo-T ribosomes not using SD sequences for either elongation or initiation. We therefore assessed the ability of Ribo-T ribosomes to use SD sequences during translation initiation. We constructed a specific dual reporter for monitoring translation efficiency in the presence of various SD sequences. We found that Ribo-T ribosomes used the SD sequence much less efficiently than wild-type ribosomes (Figure 4).

The translation initiation rate of a given gene can be modulated by the structural accessibility of the SD sequence, the thermodynamic binding potential between the SD sequence and the ASD sequence, and the exact positioning of the SD sequence relative to the start codon. Stronger SD-ASD sequence interactions are associated with a higher translation efficiency, as they stabilize the assembly of the translation initiation complex. Hockenberry *et al*. ^26^ suggested that decreasing the translation initiation rates for a large number of genes would probably lead to substantially longer doubling times. This may be the case for *E. coli* cells with Ribo-T ribosomes, which had longer doubling times and a lower translational efficiency, with a protein synthesis rate 50% that in cells with wild-type ribosomes^1^. In recent years, the role of SD sequence as the most important element governing various aspects of translation initiation (efficiency, reading frame selection, regulation) has been called into question. The existence of mRNAs devoid of the SD sequence clearly indicated that this sequence was neither necessary nor sufficient for translation initiation^27^. A local absence of RNA secondary structure has been shown to be necessary and sufficient for the initiation of SD sequence-independent translation^28^. In prokaryotic mRNAs, most of the coding sequence is highly structured and not accessible in the single-stranded form. The presence of an AUG codon in an unstructured region can define the correct translation initiation site, as recently confirmed by a genome-wide translation analysis showing the initiation of wild-type ribosomes at the correct site without the SD sequence, although the presence of the SD sequence greatly increased the efficiency of translation initiation ^29^.

The mechanism of translation initiation by Ribo-T ribosomes in the absence of the SD sequence remains unclear. However, the two subunits are linked together via helix 44 (H44) in these ribosomes^1^. This helix is known to play an important role in the fidelity of translation initiation^30^. The tethering of ribosomes may allow sufficient translation for cell viability, but it probably also has structural consequences for the geometry of H44. The ASD sequence is located at the top of H44. We therefore suggest that its orientation is modified in Ribo-T ribosomes, preventing them from using SD sequences in a normal manner during translation initiation and elongation. The elucidation of a high-resolution structure for Ribo-T ribosomes would help to resolve this question.

We also observed that Ribo-T ribosomes promoted termination less efficiently than wild-type ribosomes (Figure 5). The termination and stop codon readthrough processes are in competition every time a ribosome encounters a stop codon. The balance between these processes is displaced toward readthrough when the ribosomal A-site is occupied by a near-cognate tRNA. Not only does efficient translation termination require the coordinated action of release factors, it also depends on the conformational dynamics of the factors and the ribosome. The key conformational motions of the ribosome during termination, and throughout all phases of translation, include the rotation of ribosomal subunits relative to each other, the swiveling motion of the body and head domains of the small ribosomal subunit, the movement of the ribosomal protein L1 toward or away from the E-site tRNA, and the movement of all tRNAs^2,31,32^. The structure of Ribo-T ribosomes, in which the rRNA molecules are covalently linked, may be less dynamic, and associated with an A-site less able to bind termination factors.

These results are consistent with a previous ribosome profiling analysis in Ribo-T-expressing cells, which highlighted an increase in ribosome density at start codons and close to the 3’ ends of genes^3^. The tethering of the ribosomal subunits had mild effects on the initiation and termination/recycling steps of translation.

Our work reveals the specific behavior of tethered ribosomes and highlights the greater-than-expected flexibility of ribosomes. Undoubtedly, pushing ribosomes to their limits will also provide important information about their functioning. One question of interest concerns whether tethered ribosomes would be efficient in eukaryotes, which have a different mechanism of translation initiation not involving SD-ASD interaction.

## Acknowledgments

We would like to thank Shura Mankin for providing the strains and plasmids, sharing unpublished results and for his constructive comments about the manuscript. The English of this manuscript was corrected by Alex Edelman & Associates. This work was supported by the ANR rescue_ribosome Grant (ANR-17-CE12-0024-01).

## Bibliography

1 Orelle, C. et al. Protein synthesis by ribosomes with tethered subunits. Nature 524, 119–124, doi:10.1038/nature14862 (2015).

2 Rodnina, M. V. Translation in prokaryotes. Cold Spring Harb Perspect Biol 10, doi:10.1101/cshperspect.a032664 (2018).

3 Aleksashin, N. A. et al. Assembly and functionality of the ribosome with tethered subunits. Nat Commun 10, 930, doi:10.1038/s41467-019-08892-w (2019).

4 Rozov, A., Demeshkina, N., Westhof, E., Yusupov, M. & Yusupova, G. Structural insights into the translational infidelity mechanism. Nat Commun 6, 7251, doi:10.1038/ncomms8251 (2015).

5 Namy, O., Rousset, J. P., Napthine, S. & Brierley, I. Reprogrammed genetic decoding in cellular gene expression. Mol Cell 13, 157–168, doi:10.1016/s1097-2765(04)00031-0 (2004).

6 Namy, O. et al. Adding pyrrolysine to the *Escherichia coli* genetic code. FEBS Lett 581, 5282–5288, doi:10.1016/j.febslet.2007.10.022 (2007).

7 Stahl, G., Bidou, L., Rousset, J. P. & Cassan, M. Versatile vectors to study recoding: conservation of rules between yeast and mammalian cells. Nucleic Acids Res 23, 1557–1560, doi:10.1093/nar/23.9.1557 (1995).

8 Carlson, E. D. et al. Engineered ribosomes with tethered subunits for expanding biological function. Nat Commun 10, 3920, doi:10.1038/s41467-019-11427-y (2019).

9 Atkins, J. F., Loughran, G., Bhatt, P. R., Firth, A. E. & Baranov, P. V. Ribosomal frameshifting and transcriptional slippage: From genetic steganography and cryptography to adventitious use. Nucleic Acids Res 44, 7007–7078, doi:10.1093/nar/gkw530 (2016).

10 Polard, P., Prere, M. F., Chandler, M. & Fayet, O. Programmed translational frameshifting and initiation at an AUU codon in gene expression of bacterial insertion sequence IS911. J Mol Biol 222, 465–477, doi:10.1016/0022-2836(91)90490-w (1991).

11 Sekine, Y., Eisaki, N. & Ohtsubo, E. Translational control in production of transposase and in transposition of insertion sequence IS3. J Mol Biol 235, 1406–1420, doi:10.1006/jmbi.1994.1097 (1994).

12 Tsuchihashi, Z. & Brown, P. O. Sequence requirements for efficient translational frameshifting in the *Escherichia coli dnaX* gene and the role of an unstable interaction between tRNA(Lys) and an AAG lysine codon. Genes Dev 6, 511–519, doi:10.1101/gad.6.3.511 (1992).

13 Weiss, R. B., Dunn, D. M., Dahlberg, A. E., Atkins, J. F. & Gesteland, R. F. Reading frame switch caused by base-pair formation between the 3’ end of 16S rRNA and the mRNA during elongation of protein synthesis in *Escherichia coli*. EMBO J 7, 1503–1507 (1988).

14 Curran, J. F. & Yarus, M. Use of tRNA suppressors to probe regulation of *Escherichia coli* release factor 2. J Mol Biol 203, 75–83, doi:10.1016/0022-2836(88)90092-7 (1988).

15 Yamamoto, H. et al. 70S-scanning initiation is a novel and frequent initiation mode of ribosomal translation in bacteria. Proc Natl Acad Sci USA 113, E1180–1189, doi:10.1073/pnas.1524554113 (2016).

16 Karamyshev, A. L., Karamysheva, Z. N., Yamami, T., Ito, K. & Nakamura, Y. Transient idling of posttermination ribosomes ready to reinitiate protein synthesis. Biochimie 86, 933–938, doi:10.1016/j.biochi.2004.08.006 (2004).

17 Dabrowski, M., Bukowy-Bieryllo, Z. & Zietkiewicz, E. Translational readthrough potential of natural termination codons in eucaryotes--The impact of RNA sequence. RNA Biol 12, 950–958, doi:10.1080/15476286.2015.1068497 (2015).

18 Burke, S. A., Lo, S. L. & Krzycki, J. A. Clustered genes encoding the methyltransferases of methanogenesis from monomethylamine. J Bacteriol 180, 3432–3440 (1998).

19 Fried, S. D., Schmied, W. H., Uttamapinant, C. & Chin, J. W. Ribosome subunit stapling for orthogonal translation in *E. coli*. Angew Chem Weinheim Bergstr Ger 127, 12982–12985, doi:10.1002/ange.201506311 (2015).

20 Marquez, V., Wilson, D. N., Tate, W. P., Triana-Alonso, F. & Nierhaus, K. H. Maintaining the ribosomal reading frame: the influence of the E site during translational regulation of release factor 2. Cell 118, 45–55, doi:10.1016/j.cell.2004.06.012 (2004).

21 Liao, P. Y., Gupta, P., Petrov, A. N., Dinman, J. D. & Lee, K. H. A new kinetic model reveals the synergistic effect of E-, P- and A-sites on +1 ribosomal frameshifting. Nucleic Acids Res 36, 2619–2629, doi:10.1093/nar/gkn100 (2008).

22 Larsen, B., Wills, N. M., Gesteland, R. F. & Atkins, J. F. rRNA-mRNA base pairing stimulates a programmed −1 ribosomal frameshift. J Bacteriol 176, 6842–6851, doi:10.1128/jb.176.22.6842-6851.1994 (1994).

23 Baranov, P. V., Gesteland, R. F. & Atkins, J. F. Release factor 2 frameshifting sites in different bacteria. EMBO Rep 3, 373–377, doi:10.1093/embo-reports/kvf065 (2002).

24 Prere, M. F., Canal, I., Wills, N. M., Atkins, J. F. & Fayet, O. The interplay of mRNA stimulatory signals required for AUU-mediated initiation and programmed −1 ribosomal frameshifting in decoding of transposable element IS911. J Bacteriol 193, 2735–2744, doi:10.1128/JB.00115-11 (2011).

25 Kim, H. K. & Tinoco, I., Jr. EF-G catalyzed translocation dynamics in the presence of ribosomal frameshifting stimulatory signals. Nucleic Acids Res 45, 2865–2874, doi:10.1093/nar/gkw1020 (2017).

26 Hockenberry, A. J., Stern, A. J., Amaral, L. A. N. & Jewett, M. C. Diversity of translation initiation mechanisms across bacterial species is driven by environmental conditions and growth demands. Mol Biol Evol 35, 582–592, doi:10.1093/molbev/msx310 (2018).

27 Chang, B., Halgamuge, S. & Tang, S. L. Analysis of SD sequences in completed microbial genomes: non-SD-led genes are as common as SD-led genes. Gene 373, 90–99, doi:10.1016/j.gene.2006.01.033 (2006).

28 Scharff, L. B., Childs, L., Walther, D. & Bock, R. Local absence of secondary structure permits translation of mRNAs that lack ribosome-binding sites. PLoS Genet 7, e1002155, doi:10.1371/journal.pgen.1002155 (2011).

29 Saito, K., Green, R. & Buskirk, A. R. Translational initiation in *E. coli* occurs at the correct sites genome-wide in the absence of mRNA-rRNA base-pairing. Elife 9, doi:10.7554/eLife.55002 (2020).

30 Qin, D., Liu, Q., Devaraj, A. & Fredrick, K. Role of helix 44 of 16S rRNA in the fidelity of translation initiation. RNA 18, 485–495, doi:10.1261/rna.031203.111 (2012).

31 Korostelev, A. et al. Crystal structure of a translation termination complex formed with release factor RF2. Proc Natl Acad Sci USA 105, 19684–19689, doi:10.1073/pnas.0810953105 (2008).

32 Noller, H. F., Lancaster, L., Zhou, J. & Mohan, S. The ribosome moves: RNA mechanics and translocation. Nat Struct Mol Biol 24, 1021–1027, doi:10.1038/nsmb.3505 (2017).

